# The C-terminus and Third Cytoplasmic Loop Cooperatively Activate Mouse Melanopsin Phototransduction

**DOI:** 10.1101/2020.02.12.946129

**Authors:** J.C. Valdez-Lopez, S.T. Petr, M.P. Donohue, R.J. Bailey, M. Gebreeziabher, E.G. Cameron, J.B. Wolf, V.A. Szalai, P.R. Robinson

## Abstract

Melanopsin, an atypical vertebrate visual pigment, mediates non-image forming light responses including circadian photoentrainment and pupillary light reflexes, and contrast detection for image formation. Melanopsin-expressing intrinsically photosensitive retinal ganglion cells (ipRGCs), are characterized by sluggish activation and deactivation of their light responses. The molecular determinants of mouse melanopsin’s deactivation have been characterized (i.e. C-terminal phosphorylation and β-arrestin binding), but a detailed analysis of melanopsin’s activation is lacking. We propose that an extended 3^rd^ cytoplasmic loop is adjacent to the proximal C-terminal region of mouse melanopsin in the inactive conformation which is stabilized by ionic interaction of these two regions. This model is supported by site-directed spin labeling and electron paramagnetic resonance (EPR) spectroscopy of melanopsin, the results of which suggests a high degree of steric freedom at the 3^rd^ cytoplasmic loop, which is increased upon C-terminus truncation, supporting the idea that these two regions are close in 3-dimensional space in wild-type melanopsin. To test for a functionally critical C-terminal conformation, calcium imaging of melanopsin mutants including a proximal C-terminus truncation (at residue 365) and proline mutation of this proximal region (H377P, L380P, Y382P) delayed melanopsin’s activation rate. Mutation of all potential phosphorylation sites, including a highly conserved tyrosine residue (Y382), into alanines also delayed the activation rate. A comparison of mouse melanopsin with armadillo melanopsin—which has substitutions of various potential phosphorylation sites and a substitution of the conserved tyrosine—indicates that substitution of these potential phosphorylation sites and the tyrosine residue result in dramatically slower activation kinetics, a finding that also supports the role of phosphorylation in signaling activation. We therefore propose that melanopsin’s C-terminus is proximal to intracellular loop 3 and C-terminal phosphorylation permits the ionic interaction between these two regions, thus forming a stable structural conformation that is critical for initiating G-protein signaling.

**STATEMENT OF SIGNIFICANCE:** Melanopsin is an important visual pigment in the mammalian retina that mediates non-image forming responses such as circadian photoentrainment and pupil constriction, and supports contrast detection for image formation. In this study, we detail two critical structural features of mouse melanopsin—its 3^rd^ cytoplasmic loop and C-terminus—that are important in the activation of melanopsin’s light responses. Furthermore, we propose that these two regions directly participate in coupling mouse melanopsin to its G-protein. These findings contribute to further understanding of GPCR-G-protein coupling, and given recent findings suggesting flexibility of melanopsin signal transduction in the retina (possibly by coupling more than one G-protein type), these findings provide insight into the molecular basis of melanopsin function in the retina.

## INTRODUCTION

G-protein-coupled receptors (GPCRs) make up the largest family of integral membrane receptors and are activated by a variety of biological stimuli, including hormones, odorants, small peptides, neurotransmitters and photons (1, 2). Upon activation, GPCRs undergo a series of conformational changes that facilitate the binding and activation of intracellular heterotrimeric G-proteins, which initiate a variety of signaling responses (3, 4). Visual pigments are specialized light-detecting GPCRs that are comprised of an opsin protein covalently-attached to a chromophore, typically 11-*cis*-retinal in the mammalian retina. The absorption of a photon by 11-*cis*-retinal results in its isomerization to all-*trans*-retinal. This conformational change in the chromophore results in the activation of visual pigment and the phototransduction cascade. In the mammalian retina, three distinct classes of photoreceptor cells detect light. Rod and cone photoreceptor cells found in the outer retina express distinct specialized visual pigments that mediate photon absorption for image-forming vision. A less well-known third photoreceptor class in the inner retina is composed of a small subset of intrinsically photosensitive ganglion cells (ipRGCs) that express melanopsin, a rhabdomeric-type opsin (5, 6). In the mouse, ipRGCs are divided into six subtypes (M1-M6) that are unique in their morphology, projections into the inner plexiform layer of the retina, amount of melanopsin expressed, transcription factors, and their projections to the brain and thus, their functions (7, 8). For example, M1-ipRGCs exhibit the highest level of melanopsin expression and project to the suprachiasmic nucleus (among other brain nuclei) and thus are implicated in circadian photoentrainment (9) whereas M4-ipRGCs possess the largest soma of all ipRGC subtypes, and project to the dorsal lateral geniculate nucleus and are implicated in image formation (10). Light detected by melanopsin regulates non-image forming functions such as circadian photoentrainment, pupillary light reflex, sleep, and melatonin synthesis (9, 11, 12, 13, 14). Unlike rods and cones, melanopsin in M1-type ipRGCs signal through a Gαq-mediated pathway that leads to the opening of TRPC6/7 channels (15, 16, 17, 18). Recent studies analyzing the M4 ipRGC subtype suggest that melanopsin in these cells signals either through the Gαq transduction cascade (19) or through a cyclic-nucleotide cascade and the opening of HCN-channels (20).

Structural and molecular determinants of GPCR—G-protein complex formation have been described in both a receptor specific (for review of rhodopsin activation see 21) and systematic manner (22). However, it remains difficult to attribute specific receptor properties (e.g. amino acid sequence, polarity, charge, steric properties) to their selectivity for their cognate G-protein. The notion of a singular cognate G-protein for each receptor is also one that is challenged by evidence of receptor promiscuity to several Gα classes (23, 24) including melanopsin promiscuously activating transducin *in vitro* (25). GPCR regions important for G-protein binding selectivity include the 2^nd^ and 3^rd^ intracellular loops, and cytoplasmic extensions of transmembrane helices 5 and 6, which form critical contacts with helices 4 and 5 on Gα (26). GPCR C-termini can regulate G-protein activation as well as serve as substrates for GPCR Kinase (GRK) to mediate arrestin binding (for melanopsin C-terminal phosphorylation and deactivation, see 27, 28, 29, 30, 31, 32, 33). Structural analysis of rhodopsin—G-protein complexes reveals contacts between helix 8 on rhodopsin’s C-terminus and the C-terminal helix 5 on Gαi (34) or transducin (35), and contacts have been found on rhodopsin’s C-terminus with Gβ when in complex with Gαi (36). Additionally, rhodopsin C-terminal peptides bind and facilitate transducin signaling by enhancing phosphodiesterase activity and inhibiting transducin’s GTPase activity (37). The role of melanopsin’s cytoplasmic loops or C-terminus in stimulating G-protein activity remain to be determined.

In this study, we aimed to determine the critical cytoplasmic domains on mouse melanopsin that contribute to phototransduction activation. Melanopsin-mediated behaviors are diverse and are both image- and non-image-forming. Additionally, ipRGC light responses are sluggish (single flash responses can persist across minutes) (38) compared to rapid photoresponses observed in rod and cone photoreceptors (these cells can respond in a precise manner to millisecond-scale light flashes). Melanopsin Gαq signal transduction has been shown to be re-purposed to modulate leak potassium channels (rather than activating phospholipase C) or could possibly couple to another G-protein to stimulate cyclic nucleotide signal transduction, in M4-type ipRGCs (19, 20). Within the M1-type population, there is a striking heterogeneity in the light responses, where there is a mixture of cells that optimally respond to certain irradiances or respond in a linear fashion in response to increasing irradiance, thus allowing for this M1 population to encode for a breadth of light intensities (39). Additionally, melanopsin phosphorylation by Protein Kinase A, likely on the intracellular loops, can attenuate light responses in a dopamine and cyclic AMP-dependent manner (28). It is therefore critical to examine melanopsin—G-protein complex formation at a structural level to shed light on the molecular determinants underlying the functional diversity observed in melanopsin signal transduction. Based on amino acid sequence analysis and homology modeling using *Todarodes pacificus* rhodopsin as a template, melanopsin is predicted to have extended 5^th^ and 6^th^ transmembrane helices and also a uniquely long C-terminus (Figure 1 for 2D schematic of amino acids, Figure 2 for 3D figures), which is 171 amino acids long (residues T350-L521). Given the functional significance of these regions in other GPCRs, we synthesized a series of mutants designed to test if these regions contribute to phototransduction activation. Our findings provide a detailed examination of the structural basis of melanopsin activation and, furthermore, provide a more robust foundation to examine ipRGC activation and structure—function relationships of other related visual pigments.

**Figure 1:**
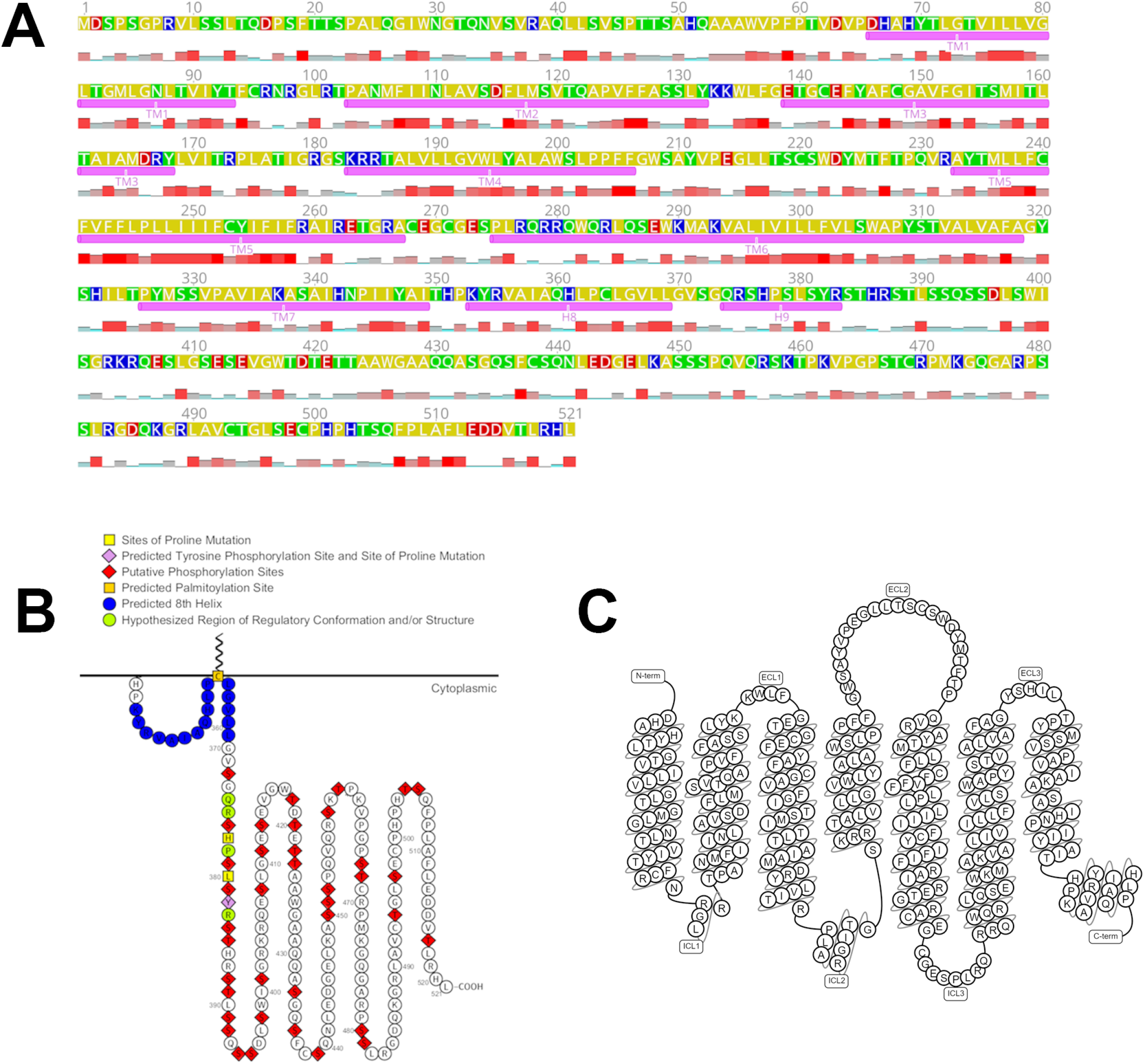
Mouse melanopsin’s amino acid sequence. Mouse melanopsin is 521 amino acids long, with a long C-terminus domain that is 171 amino acids long. (**A**) Transmembrane and cytoplasmic helices are annotated below the amino acid sequence. Amino acid property is denoted by the color of the residue: Yellow: Nonpolar, Green: Polar, non-charged, Blue: Positively charged, Red: Negatively charged. Hydrophobicity at each residue is plotted below the amino acid sequence, taller and red bars indicate a high level of hydrophobicity at that position. Identities of residues possessing secondary structures are based on prediction from the homology mode generated (See Figure 2). Figure generated using Geneious software (63). (**B**) 2-dimensional schematic of mouse melanopsin C-terminus depicting functionally significant residues. Figure made using Protter software (64). (**C**) 2D snakeplot of mouse melanopsin’s secondary structure. Figure from GPCRdb, using SWISSPROT for secondary structure prediction (65). Note: the amino acids labeled as transmembrane helices in (A), not (C), correspond to identities of the amino acids in the homology model in Figure 2.

## EXPERIMENTAL PROCEDURES

### Homology modeling of mouse melanopsin

Mouse melanopsin amino acid sequence (Uniprot ID: Q9QXZ9-1; OPN4L) was used to produce a homology model using the online protein structure service LOMETS, which ranks models derived from 11 servers using a Z-score as a metric of confidence for each model (40). The melanopsin structural model that was generated used *Todarodes pacificus* rhodopsin (PDB ID: 2ZIY) (48) as the template, which was the highest ranked model and was produced using the HHpred threading program (41). To model the active conformation of mouse melanopsin, the LOMETS-generated homology model was aligned to the crystal structure of the β_2_-adrenergic receptor in complex with Gαs (PDB ID: 3SN6) (46) and cryo-EM structure of rhodopsin in complex with Gαi (PDB ID: 6CMO) (34) in PyMol (Schrödinger Inc, New York, NY) using the cealign tool, and predicted melanopsin helical movements were modeled using the positions of the transmembrane helices of the β_2_-adrenergic receptor or rhodopsin structures. Surface electrostatic potential of melanopsin was calculated using vacuum electrostatics function on PyMol.

### Synthesis of Melanopsin Mutants

All mutant genes were constructed using the mouse melanopsin coding sequence (NCBI accession: NM_013887.2) cloned in the mammalian expression plasmid PMT3 (42). Melanopsin Δ365, H377A L380A Y382A, phosphonull + Y382S, melanopsin ICL3 Null, melanopsin 280-285 alanine, armadillo melanopsin constructs, and Mouse 1D4-Gαq (*Mus musculus Gnaq* with N-terminal 1D4 tag—amino acid sequence TETSQVAPA, corresponding to the last 9 amino acids of bovine rhodopsin, NCBI accession: NM_008139.5) were synthesized by cassette synthesis using synthetically-synthesized Gblock gene fragments (IDT, Coralville, IA). Melanopsin C268 was constructed using Gibson assembly (43) of synthetically-synthesized Strings DNA fragments (ThermoFisher Scientific, Waltham, MA) to generate mutagenesis of residues C95, C271, C438, C469, C493, C499 to alanine residues to eliminate non-specific spin label attachment. These constructs for EPR spectroscopy were also C-terminally tagged with the 1D4 epitope to facilitate immunoprecipitation when expressed heterologously. Melanopsin C268 Δ365 was synthesized through cassette synthesis using melanopsin C268 as a PCR template and with a reverse primer to amplify from the start of the gene to the end of the codon encoding for L365. Melanopsin H377P L380P Y382P, phosphonull + Y382A, and phosphonull + Y382F were generated using Quickchange-based site-directed mutagenesis (44). All plasmids were sequence verified through Sanger sequencing (Genewiz, South Plainfield, NJ).

### Cell Culture and Transfection of HEK293 Cells

HEK293 cells were grown in a monolayer on 10 cm culture dishes in Dulbecco’s Modified Eagle Medium (DMEM) (ThermoFisher Scientific, Waltham, MA) supplemented with fetal bovine serum (10% v/v) and antibiotic-antimycotic (1% v/v) (ThermoFisher Scientific, Waltham, MA) in a humid CO_2_ incubator at 37 °C. Cells were passaged by disassociation using (0.25% w/v) trypsin-EDTA (ThermoFisher Scientific, Waltham, MA) and seeded to a density of up to 5,000,000 cells per 10 cm dish and 240,000 cells per well in a 6 well dish. Transfections were done using Turbofect Transfection Reagent (ThermoFisher Scientific, Waltham, MA) as per manufacturer’s protocol. Briefly, 4 or 10 μg DNA (6-well and 10 cm dish, respectively) was diluted in non-supplemented DMEM, and then the transfection reagent was added to the mixture. Following a 20 min incubation at room temperature, the transfection mixture was added to the cells, and the cells were incubated in a humid CO_2_ incubator for 24 h to 48 h.

### Calcium Imaging of Melanopsin-Transfected HEK293 Cells

24 h after transfection, HEK293 cells were tryspin-disassociated (0.25% w/v trypsin-EDTA) from the plates and seeded to 96-well plates at a density of 100,000 cells per well. The 96-well plate is incubated overnight in a humid CO_2_ incubator in a dark room for dark adaptation. The cells are then incubated with equivalent volume of Fluo-4 AM Direct Calcium Assay Dye (ThermoFisher Scientific, Waltham, MA) supplemented with 20 mM probenecid and 20 μM 9-*cis*-retinal (Sigma Aldrich), and are incubated for 1 hr in a humid CO_2_ incubator. Calcium kinetics were then measured on a TECAN Infinite M200 (TECAN Trading AG, Morrisville, NC) by exciting the sample at 487 nm and recording the emission fluorescence at 516 nm at a rate of 1 Hz. The plate reader monochromator has a bandwidth of <9 nm for excitation light. Every melanopsin construct in each transfection had six replicates.

### Protein expression and preparation for EPR spectroscopy

Melanopsin C268 and C268 Δ365 constructs were transfected in HEK293 cells in 60-80 10 cm plate batches, harvested 48 hr later, and stored at −80 °C. Frozen cell pellets were then thawed on ice then immediately resuspended in PBS with 40 μM 9-*cis*-retinal (Sigma-Aldrich, St. Louis, MO) and incubated for at least an hour at 4 °C to reconstitute the visual pigment. The sample was centrifuged at max speed on a clinical centrifuge and the cell pellet was then resuspended in solubilization buffer composed of 0.1% (w/v) *n*-dodecyl-β-D-maltoside, 3 mM MgCl2, 140 mM NaCl, 1 mM phenylmethylsulfonyl fluoride (PMSF), and 50 mM HEPES, pH 7.5. After a minimum of 1 hr on a tube rotator at 4 °C, the solubilized cell lysate was centrifuged at max speed on a clinical centrifuge and the supernatant was collected. The solubilized protein was then incubated with Sepharose-4B resin (GE Healthcare Life Sciences, Pittsburgh, PA) conjugated with monoclonal α-1D4 antibody (provided generously by Prof. Daniel Oprian) for at least an hour at 4 °C. The 1D4-resin was then washed by centrifuging 30 secs at 500 x g, removing the supernatant, and resuspending the resin with solubilization buffer at least 10 times. After the final wash, the 1D4-resin was resuspended in solubilization buffer containing 2 mM MTSL to attach the spin label at C268 for EPR spectroscopy, and was incubated overnight on a tube rotator at 4 °C. The sample was then washed by centrifuging 30 secs at 500 x g, removing the supernatant, and resuspending the resin with solubilization buffer at least 5 times. MTSL spin-labeled melanopsin was then eluted by incubating the 1D4-resin in solubilization buffer containing 50 μM 1D4 peptide (amino acid sequence TETSQVAPA). Eluate was centrifuged using Amicon Ultra 10K centrifugal filters (Millipore Sigma, Burlington, MA) to concentrate the sample and filter out the eluting peptide.

### EPR Spectroscopy of single spin-labeled melanopsin

CW EPR spectra at X-band (approximately 9.8 GHz) were recorded using a commercial EPR spectrometer equipped with a commercial microwave cavity in perpendicular mode. Samples were drawn, by capillary action, into calibrated 25 μL disposable borosilicate glass micropipettes sealed with a hematocrit tube sealing putty, and subsequently inserted into a critically coupled EPR cavity at ambient temperature. Spectra (88 averages) were recorded using 0.1 mT modulation amplitude at 100 kHz, with an incident microwave power of 1.5mW. Simulations of EPR data was done using EasySpin (45).

### Calculation of Melanopsin Activation Rates

The activation phases of calcium imaging assay data (corresponding to the part of the data prior to the peak fluorescence level) for all melanopsin constructs was fitted to a one-phase association function using GraphPad Prism software (GraphPad Software, Inc., San Diego, CA). The following function was used:

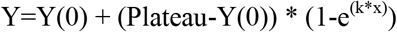

Where Y(0) is the value at t=0, Plateau is the peak activation value, and k is the rate constant. Data across multiple transfections (six replicates per transfection) were pooled together and averaged, and the activation rate of the averaged data was calculated. Standard error of the mean of the deactivation rates of all melanopsin constructs and statistical analysis were calculated using GraphPad software. Statistical significance was determined by performing unpaired t-tests of all mutant melanopsin constructs with respect to wild-type melanopsin, or between specific melanopsin constructs of interest.

## RESULTS

### Three-dimensional homology modeling of mouse melanopsin predicts that an extended 3^rd^ cytoplasmic loop and the proximal C-terminus mediate G-protein binding

The structural basis of melanopsin activation is not well understood; specifically, the unique molecular mechanisms underlying its capability to couple its cognate G-protein have not been elucidated in detail. To shed light on this problem, we constructed a homology model of mouse melanopsin using squid rhodopsin as a template. The generated homology model had 68 % of melanopsin’s amino acid sequence aligned to the template (Supplemental Figure 1), and among these amino acids, they shared 37 % sequence identity to the squid (*Todarodes pacificus*) rhodopsin template. The sequence coverage of our model includes the entirety of the seven-transmembrane helical domain and the proximal portion of the C-terminus, up to S398. In the inactive state (Figure 2A), the model predicts that melanopsin forms a seven-transmembrane visual pigment with an extended 3^rd^ intracellular loop, which is comprised of cytoplasmic extensions of transmembrane helices 5 and 6. In the active state, which we modeled based on helical movements of β_2_-adrenergic receptor (46) and rhodopsin (34), melanopsin’s 5^th^ and 6^th^ transmembrane helices are predicted to swing away from the C-terminus (Figure 2A), thereby making way for the attachment of G-protein to the newly formed binding pocket (Figure 2B-K). The generated homology model also predicts a C-terminal 9^th^ helix (Figure 2A), but further experimental data is needed to test the existence of this structure in melanopsin, but the data presented in this study strongly support the predicted C-terminal conformation shown in the homology model.

**Figure 2:**
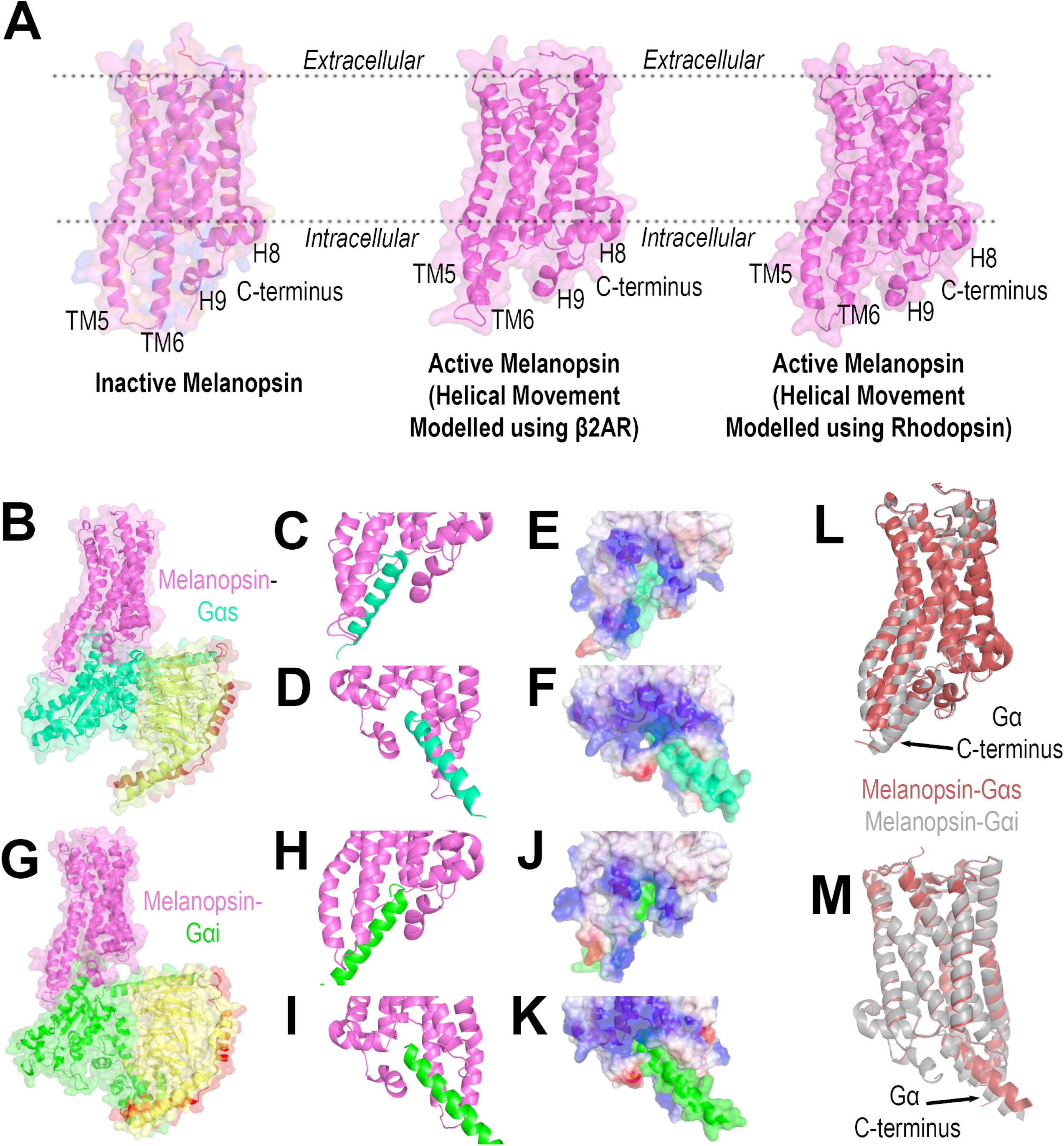
Homology modeling of mouse melanopsin. (**A**) Inactive and active conformations of mouse melanopsin, generated using squid (*Todarodes pacificus*) rhodopsin as a template, and using β_2_ adrenergic receptor or rhodopsin to model helical movements in the active melanopsin models. (**B**) Melanopsin in complex with Gαs. Purple: melanopsin, Green: Gα, Yellow: Gβ, Dark Red: Gγ. (**C & D**) Zoomed in views of mouse melanopsin’s (purple) cytoplasmic domains when in complex with Gαs C-terminal peptide (green). (**E & F**) Zoomed in views depicted in (C) and (D), depicted as a space-filling model with melanopsin’s electrostatic potentials plotted on the surface (Blue: positively charged, Red: negatively charged, White: Neutral). (**G**) Melanopsin in complex with Gαi. Purple: melanopsin, Green: Gα, Yellow: Gβ, Dark Red: Gγ. (**H & I**) Zoomed in views of mouse melanopsin’s (purple) cytoplasmic domains when in complex with Gαi C-terminal peptide (green). (**J & K**) Zoomed in views depicted in (H) and (I), depicted as a space-filling model with melanopsin’s electrostatic potentials plotted on the surface (Blue: positively charged, Red: negatively charged, White: Neutral). (**L & M**) Overlay of Gαs and Gαi complexed melanopsin structures.

The crystal structures obtained for squid rhodopsin reveal contacts between intracellular loop 3 (specifically, the cytoplasmic extension of transmembrane helix 6) and its C-terminal 9^th^ helix (47, 48). Additionally, the presence of positively charged residues on squid rhodopsin’s 3^rd^ intracellular loop and negatively charged residues on the C-terminus, may facilitate the formation of contacts between these regions, or potentially to the Gαq C-terminus. Additionally, while most GPCRs display promiscuous binding to several Gα-subtypes, several molecular features on the GPCR determine the selectivity for cognate G-proteins. Systematic analysis of GPCRs suggest that selectivity to the Gαq/11 family is determined primarily by the amino acid properties of the receptor’s C-terminus, specifically through enrichment of non-aromatic residues (22). Interestingly, the lengths of either the C-terminus or the 3^rd^ intracellular loop are not positively correlated with selectivity to Gαq/11-subtypes and are only selectivity features in the binding of Gα12/13 (22). Surface electrostatic analysis of our homology model of melanopsin suggests an enrichment of positively charged residues on the intracellular loops, particularly intracellular loop 3 (Figure 2E & 2J) and 2 (Figure 2F & 2K). The proximal region of melanopsin’s C-terminus is also positively charged (Figure 2E & 2J), and we predict it will make contacts with positively-charged residues on transmembrane helix 6 in the inactive conformation (Figure 2A). Thus, the ionic contacts between the C-terminus and 3^rd^ intracellular loop would require amino acid modification (namely, C-terminal phosphorylation) to provide the necessary charge for these ionic contacts. Comparison of both our active state melanopsin models suggests that melanopsin would have to undergo different helical movements, specifically a greater outward swing of transmembrane helix 6 when melanopsin is modeled with Gαs (Figure 2L & 2M). Systematic analysis of GPCRs suggests no positive correlation with negative surface charge at these critical cytoplasmic domains on the receptor and their capability to couple Gαq (22). However, we propose that the amino acid ionic charges and the conformation of melanopsin’s cytoplasmic domains play critical roles in its ability to couple to G-protein(s) and initiate ipRGC phototransduction.

### Electron paramagnetic resonance (EPR) spectroscopy of single spin-labeled mouse melanopsin supports the proposed C-terminus—intracellular loop 3 arrangement

To test our model that predicted the C-terminus and cytoplasmic loop, we synthesized a mutant melanopsin for EPR spectroscopy analysis, Melanopsin C268. All solvent-accessible, non-specific cysteines were also mutated to alanine residues with the exception of C364, a predicted palmitoylation site, and residues C142 and C220, which are predicted sites of a disulfide bridge, a common molecular feature of opsins necessary for proper folding and retention of covalently-linked retinal molecule (49, 50). This mutant was designed for single labeling using the nitroxide spin label MTSL (2,2,5,5-tetramethyl-1-oxyl-3-methyl methanethiosulfonate) at C268 on intracellular loop 3 to analyze the steric freedom at this position in the presence (Melanopsin C268) or absence (Melanopsin C268 Δ365) of the C-terminus.

The continuous wave (CW) EPR spectrum of single spin-labeled, dark-adapted melanopsin C268 (Figure 3A) in 0.1 % (w/v) *n*-dodecyl-β-D-maltoside (DM) presents a sharper spectral lineshape compared to similar EPR measurements on rhodopsin (51, 52), a finding that indicates a high degree of rotational mobility of the MTSL spin label at this position. In addition to the qualitative observation of a sharper spectral lineshape, mathematical fitting of the EPR spectrum of single spin-labeled melanopsin C268 to extract a rotational correlation time (*τ*_corr_) of the spin label supports the higher degree of steric freedom at this region compared to rhodopsin. The calculated *τ*_corr_ of melanopsin C268 was 9.36 ns (Supplemental Figure 2), which is lower than *τ*_corr_ of both inactive and active forms of rhodopsin (77 ns and 26 ns, respectively) when spin-labeled at the cytoplasmic end of transmembrane helix 6 (V250C) (53). One possible explanation for the increased spin label rotational mobility in the C268-labeled melanopsin is that the 3^rd^ intracellular loop is extended into the cytoplasm, as predicted by the structural model. The broader lineshape and longer rotational correlation times for rhodopsin (51, 52, 54) suggest that the spin label’s rotational mobility (MTSL attached at the cytoplasmic face of transmembrane helices) is hindered due to closer proximity to the DM micelle compared to melanopsin’s spin label. The CW EPR spectrum of single spin-labeled, dark-adapted C-terminal truncated melanopsin (Melanopsin C268 Δ365) in DM (Figure 3B) possesses an even sharper spectral lineshape than full-length melanopsin and resembles the lineshape of the spectrum observed when measuring free MTSL in 0.1 % (w/v) DM solution, without protein (Figure 3C). Together, these data indicate a higher degree of MTSL rotational mobility at the 3^rd^ intracellular loop, which we attribute to the elimination of any steric hinderance caused by the C-terminus consistent with the C-terminal conformation proposed in Figure 2.

**Figure 3:**
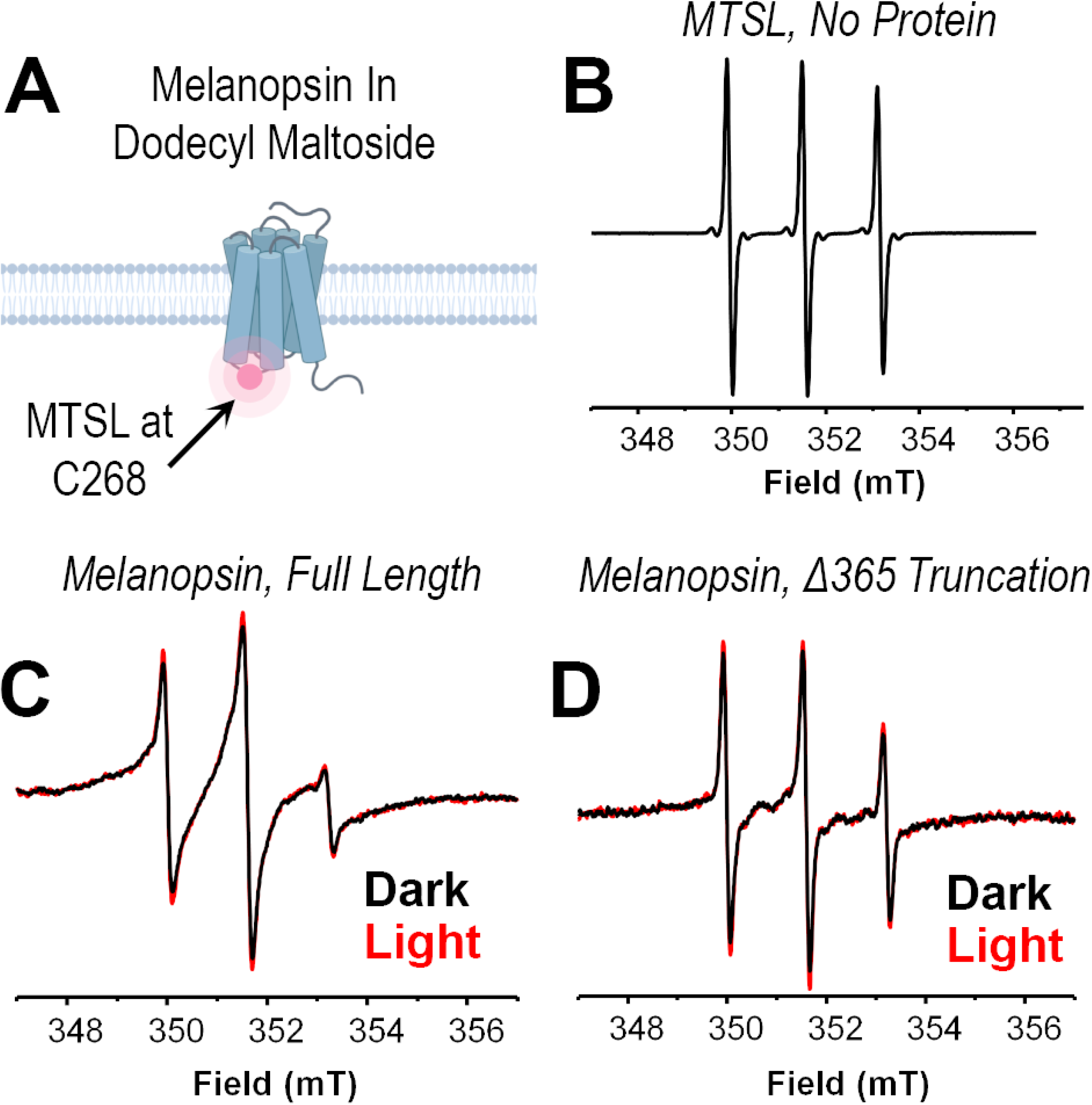
Electron paramagnetic resonance (EPR) spectroscopy of site-directed spin-labeled mouse melanopsin in n-dodecyl-β-D-maltoside. (**A**) Cartoon schematic of the EPR measurement of melanopsin. Affinity-purified melanopsin in dodecyl maltoside solution was site-directed spin-labeled at C268, on the 3^rd^ cytoplasmic loop. (**B**) EPR spectrum of MTSL in solution with no protein. (**C & D**) EPR spectra of mouse melanopsin, MTSL-conjugated at C268. EPR spectra were recorded in darkness under dim red-light illumination at room temperature. (**C**) EPR spectrum of full length melanopsin C268 in darkness (black trace) and after 30 sec light illumination (red trace). (**D**) EPR spectrum of C-terminally truncated melanopsin C268 in darkness (black trace) and after 30 sec light illumination (red trace).

After measurement of the EPR spectra of full length and truncated Melanopsin C268, the samples were exposed to 30 s of 300 W white light to test for differences in the spectra upon light-induced conformational change of the visual pigment. Comparison of the dark and light EPR spectra indicates minimal differences between the two spectra of each sample (Figure 3A & 3B), indicating that the rotational mobility of the spin label at position C268 is minimally affected by light-induced conformational change.

### The proximal region of mouse melanopsin’s C-terminus is necessary for rapid phototransduction activation

To test the hypothesis that melanopsin’s C-terminus regulates signaling initiation, we synthesized melanopsin C-terminal mutants (positions in Figure 1B) aimed to disrupt or ablate C-terminal structure or configuration in its proximal region (prior to residue D396). To test for potential C-terminal regulation of phototransduction activation, we synthesized a mouse melanopsin C-terminus truncation mutant, Melanopsin Δ365, which is truncated at residue L365, and eliminates the C-terminal region predicted to form part of the G-protein binding pocket in our structural model. Calcium imaging of this mutant reveals delayed activation kinetics (Figure 4A) compared to wildtype melanopsin, supporting a functional role of this C-terminal region in mediating signaling activation. Previous studies that examined melanopsin C-terminal truncations (29) suggest that the distal C-terminus (after D396) doesn’t contribute to the production of rapid signaling activation and can potentially sterically hinder the G-protein from accessing the cytoplasmic side of the receptor (32). To disrupt the predicted proximal C-terminal conformation, we introduced proline mutations (H377P L380P Y382P) in the critical region predicted to work synergistically with the cytoplasmic loops to regulate phototransduction activation. These proline mutations introduced backbone rigidity and thus will disrupt any important conformations or potential secondary structures found in the wild-type protein (55). Calcium imaging of this triple proline mutant (Figure 4B) reveals that it has delayed activation kinetics compared to wildtype melanopsin, indicating this region has a functionally significant conformation, as suggested by the structural modeling. Mutation of those three residues produces the most pronounced reduction in activation rate, as pairwise proline melanopsin mutants H377P L380P and L380P Y382P show slight, but not statistically significant reductions in activation rate, while the H377P Y382P and H377P L380P Y382P show significant activation rate reduction (Supplemental Figure 3). To test that the activation defect observed in the proline mutant is attributable to a proline-induced kink or disruption in structure, we synthesized and tested another mutant of melanopsin with alanine mutations at the same sites of interest in the proline mutant (H377A L380A Y382A). We predicted that these mutations would not disturb any functionally important C-terminal structure or conformation, unlike the proline mutations. Calcium imaging of this mutant shows no reduction in the activation rate, suggesting that the proline mutant’s defect is indeed due to a disruption of C-terminal structure or conformation.

**Figure 4:**
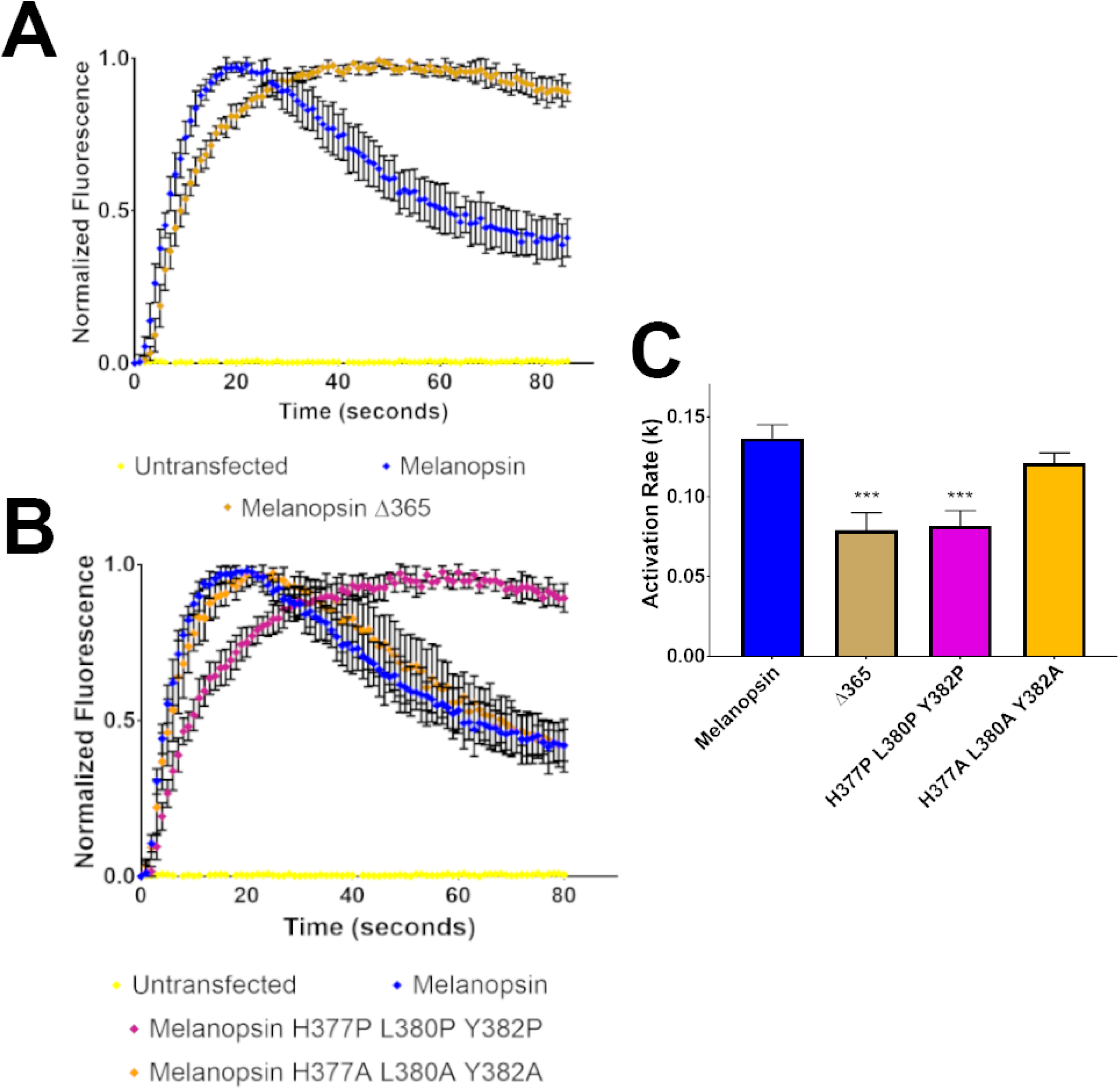
Calcium imaging of melanopsin C-terminal mutants. (**A**) Calcium imaging of HEK293 cells expressing melanopsin Δ365, a C-terminus mutant truncated at residue L365 and (**B**) calcium imaging of melanopsin H377P L380P Y382P, a C-terminus mutant with point mutations designed to disrupt the critical conformation at this region. (**C**) Calculated activation rates of melanopsin constructs tested in these experiments. All error bars represent S.E.M. of three independent transfections. Statistical significance tested by using Students t-test, *, **, ***, **** represent P-values <0.05, 0.01, 0.001, and 0.0001, respectively. All constructs compared to wild-type melanopsin’s rate, and statistical significance is indicated over individual bars.

C-terminal palmitoylation anchors and stabilizes GPCR C-termini (56) and in rhodopsin, it regulates its proper expression and stability in the membrane (57, 58). Mutation of mouse melanopsin’s predicted palmitoylation site from a cysteine to serine residue and calcium imaging of this mutant, melanopsin C364S, reveals a very modest increase in this mutant’s time to reach peak activation, and a slower deactivation compared to wild-type melanopsin (Supplemental Figure 4). These data suggest that C364, if palmitoylated, might stabilize the proximal C-terminus for signal transduction, possibly in a cooperative manner with the critical C-terminal conformation downstream of this site. Taken together, we propose that the proximal region of the C-terminus, between residues L365 and D396, holds functional significance that is dependent on an important structural conformation of these residues.

### Proximal C-terminal phosphorylation regulates melanopsin phototransduction activation kinetics

Mouse melanopsin’s C-terminus possesses 38 serine and threonine residues (Figure 1B) that may serve as potential phosphorylation sites. It is well established that melanopsin C-terminal phosphorylation is critical for signaling deactivation and the lifetime of melanopsin-driven behaviors such as pupil constriction and jet-lag photoentrainment (28, 29, 31, 32). Given the enrichment of serine and threonine residues at the proximal region (from residues H351-T385) of mouse melanopsin’s C-terminus, we tested if phosphorylation of these sites contributes to regulation of phototransduction activation, in addition to deactivation. To test this idea, we performed calcium imaging of transiently-transfected HEK293 cells expressing melanopsin C-terminal phosphorylation mutants: phosphonull melanopsin (28, 31), a mutant with all 38 C-terminal serine and threonine residues mutated to alanine residues, and phosphonull + Y382A melanopsin, a mutant with an evolutionarily conserved tyrosine residue (Figure 5A) mutated to an alanine residue in addition to the phosphonull C-terminus mutations. Calcium imaging of these mutants (Figure 5B) suggests that melanopsin C-terminal phosphorylation contributes to signaling activation, due to both mutants displaying slower rates of activation (Figure 5C) compared to wildtype melanopsin. Phosphonull + Y382A melanopsin displays a slower rate of activation than phosphonull melanopsin, similar to melanopsin Δ365 and melanopsin H377P L380P Y382P mutants. This implies that Y382, an evolutionarily conserved residue in mammalian melanopsins, holds key significance for regulating the kinetics of signaling activation.

**Figure 5:**
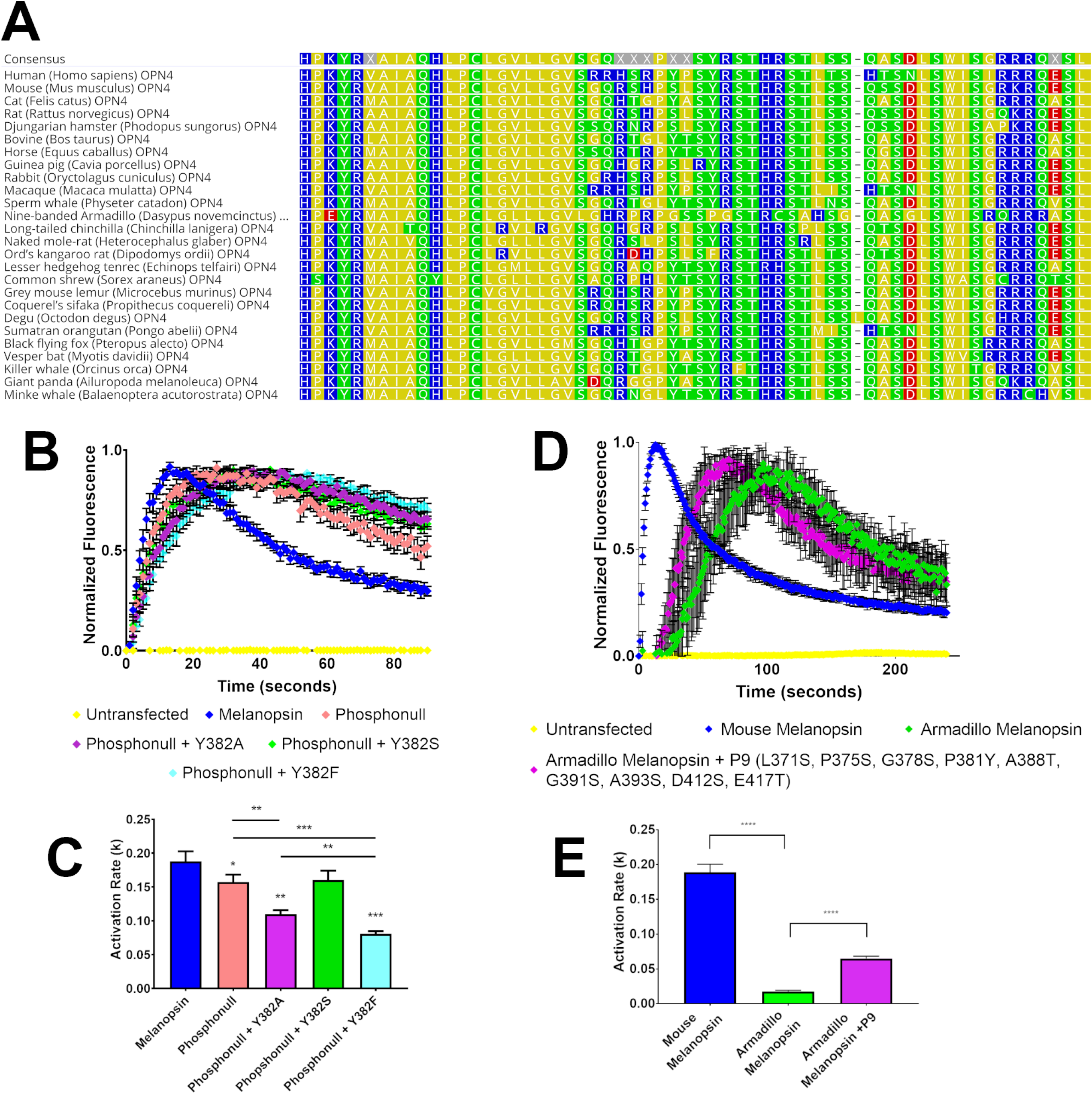
Calcium imaging of melanopsin C-terminal phosphorylation mutants. (**A**) Amino acid alignment of mammalian melanopsins focused on the proximal C-termini. Arrow depicts the conserved tyrosine, Y382 in mouse melanopsin. Alignment generated using Geneious software (Kearse et al, 2012). (**B**) Calcium imaging of HEK293 cells expressing melanopsin C-terminal phosphorylation mutants (**C**) Calculated activation rates of melanopsin constructs depicted in (B). (**D**) Calcium imaging of HEK293 cells expressing mouse and armadillo melanopsins. (**E**) Calculated activation rates of melanopsin constructs depicted in (D). All error bars represent S.E.M. of three independent transfections. Statistical significance tested by using Students t-test, *, **, ***, **** represent P-values <0.05, 0.01, 0.001, and 0.0001, respectively. All constructs compared to wild-type melanopsin’s rate, and statistical significance is indicated over individual bars. Additional statistical analyses between constructs are indicated with the lines above the appropriate bars.

To elucidate the mechanism of this tyrosine residue, specifically, its potential phosphorylation to facilitate ionic interaction with intracellular loop 3, we synthesized additional point mutations at that position, specifically, phosphonull + Y382S and phosphonull + Y382F (Figure 5B). By mutating Y382 to serine and phenylalanine residues, we aimed to test if potential phosphorylation at this position (by means of the phosphorylatable residue, serine) contributes to activation kinetics. Specifically, if the elimination of phosphorylation, but retention of the atomic mass (phenylalanine has a similar aromatic structure as tyrosine but lacks the hydroxyl group for phosphorylation modification) produces a similar defect as phosphonull + Y382A, then we can conclude that the addition of a phosphate group is important. Calcium imaging of these mutants reveals that phosphonull + Y382S has a higher rate of activation compared to phosphonull + Y382A, and has a similar rate as phosphonull and is not statistically lower than wild-type melanopsin’s rate of activation (Figure 5C). This supports the importance of potential phosphorylation at this position. Analysis of the activation rate of phosphonull + Y382F shows that it has slow activation kinetics, slower than phosphonull + Y382A (Figure 5C), supporting the conclusion that charge and not atomic size is important in this region of melanopsin.

To further test the evolutionary basis of C-terminal phosphorylation and the proposed functional tyrosine, Y382, in signaling activation, we transiently transfected armadillo melanopsin in HEK293 cells and tested its signaling kinetics using calcium imaging. Armadillo melanopsin possesses a shorter C-terminus than mouse melanopsin, and it has fewer C-terminal serine and threonine residues than mouse melanopsin (Armadillo melanopsin has 14 potential phosphorylation sites compared to 38 in mouse melanopsin, Supplemental Figure 5). Additionally, armadillo melanopsin has a substitution of the evolutionarily conserved tyrosine residue (P381 in armadillo melanopsin, Figure 5A), which may potentially directly impact the activation kinetics. Calcium imaging of mouse and armadillo melanopsins (Figure 5D) indicates a striking difference in signaling kinetics, most notably, armadillo melanopsin’s activation kinetics are significantly slower than mouse melanopsin (Figure 5E) and displays a brief time period of inactivity before phototransduction onset. This suggests that the lack of C-terminal serine and threonine residues, particularly the proximal residues, and the lack of the tyrosine residue in armadillo melanopsin directly affects its capability to activate phototransduction. Furthermore, point mutations of armadillo melanopsin to mimic mouse melanopsin residues (L371S P375S G378S P381Y A388T G391S A393S D412S E417T) resulted in faster activation kinetics (Figure 5D & 5E) than wild-type armadillo melanopsin, further supporting the importance of C-terminal phosphorylation in producing rapid signaling activation.

### Positively-charged residues on mouse melanopsin’s 3^rd^ intracellular loop play a critical role in G-protein activation

The functional importance of GPCR intracellular loop domains in the coupling and activation of its cognate G-protein has been established in many receptors. We wanted to test what molecular features of melanopsin’s cytoplasmic loops confer its capability to activate G-protein signaling. We focused on the 3^rd^ intracellular loop due to its extended nature and predicted interaction with the C-terminus. In addition, melanopsin’s 3^rd^ intracellular loop is rich with positively-charged residues, and we wanted to test if this property has any role in phototransduction activation. First, we synthesized a melanopsin mutant where amino acids 280-285 (R280 Q281 W282 Q283 R284 L285) on the cytoplasmic extension of transmembrane helix 6 were mutated to alanine residues (Melanopsin 280-285 Alanine, Figure 6A) to disrupt any hydrogen-bonding or ionic contacts with the C-terminus, as predicted by our modeling. Calcium imaging of this mutant reveals sluggish activation kinetics (Figure 6D), similar to our C-terminal activation mutants (Figure 4). This data supports the model of functionally significant contacts between intracellular loop 3 and the C-terminus. To test for any functional significance of the negative charge localized at the 3^rd^ cytoplasmic loop, we mutated all positively-charged residues on the 3^rd^ intracellular loop (Figure 6B) to alanine residues (Melanopsin R259A R262A R266A R277A R279A R280A R284A K390A K293A or Melanopsin ICL3 Null, Figure 6C) to alter the loop’s surface charge closer to neutral. Calcium imaging of this mutant shows a striking inability to activate phototransduction (Figure 6D), suggesting that the negative charge at the 3^rd^ intracellular loop is critical for melanopsin to couple to G-protein in any capacity, thus we propose this feature as a contributor to melanopsin’s capability to couple to Gα, or potentially also its selectivity for its cognate G-protein.

**Figure 6:**
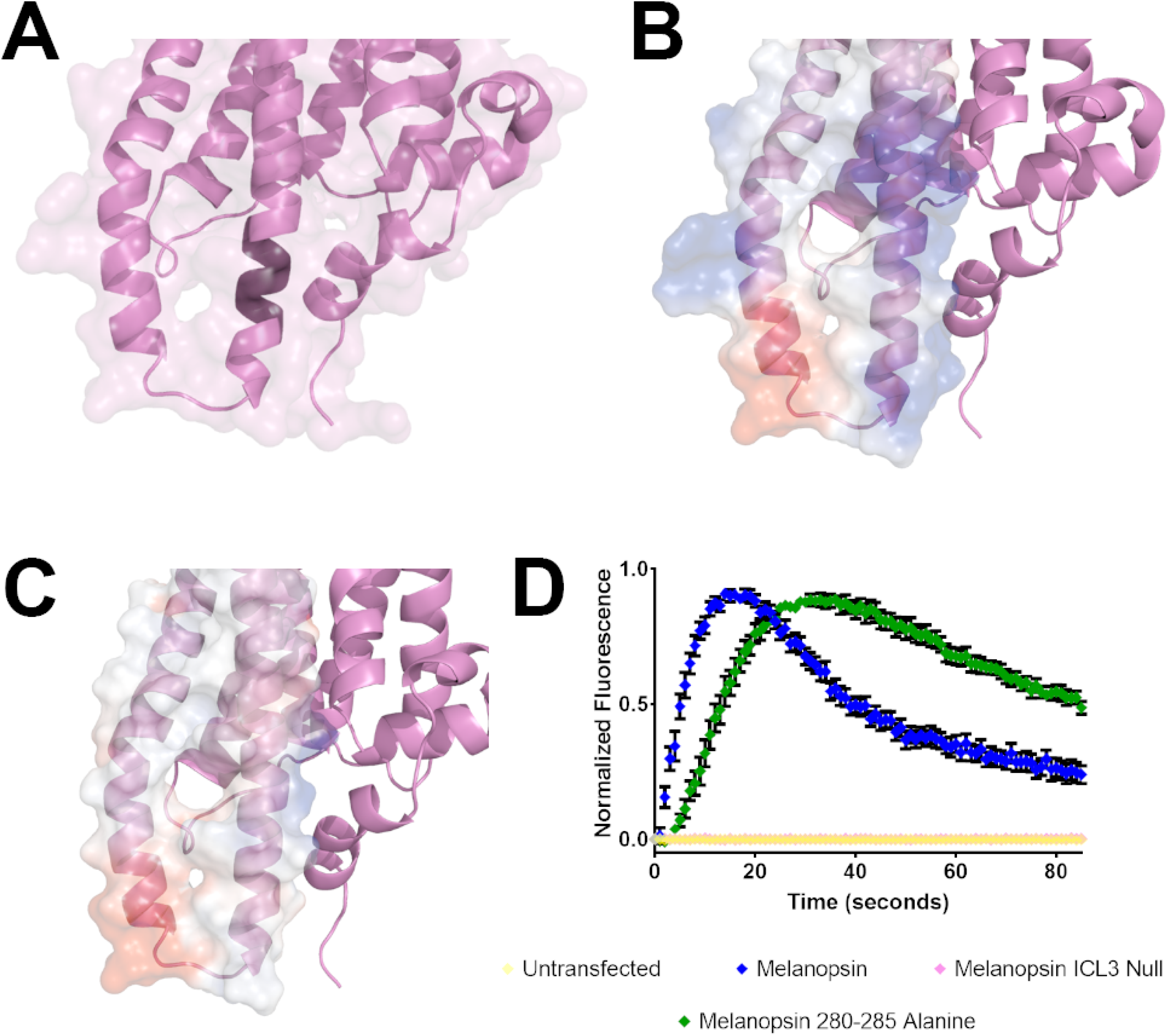
Calcium imaging of intracellular loop 3 mutants. (**A**) Homology model of melanopsin depicting the region of mutations (darker purple color on the cytoplasmic extension of transmembrane helix 6) in the mutant, melanopsin 280-285 Alanine, where amino acids 280-285 are mutated to alanine residues. (**B**) Homology model depicting the electrostatic charge of the 3^rd^ cytoplasmic loop in wildtype melanopsin and (**C**) the mutant melanopsin ICL3 Null, where all positively-charged residues were mutated to alanines to neutralize the charge of this region (Blue: positively charged, Red: negatively charged, White: Neutral). (**D**) Calcium imaging of melanopsin and the mutants depicted in (A) and (C). Error bars represent S.E.M. of three independent transfections

## DISCUSSION

The diversity of molecular features and amino acid properties in the cytoplasmic regions of GPCRs complicates the prediction of receptor-G-protein binding properties in an unstudied receptor. It is evident in many class A GPCRs, including the prototypical receptor and visual pigment rhodopsin, that cytoplasmic domains including the cytoplasmic loops and the C-terminus are critical for G-protein binding and selectivity. Melanopsin-expressing ipRGCs in the mammalian retina regulate a diversity of non-visual and visual functions and functional heterogeneity is observed across ipRGC subtypes and even within the same subtypes. In this study, we propose our model of melanopsin consisting of a functionally critical cytoplasmic conformation consisting of extended transmembrane helices 5 and 6 that form a 3^rd^ cytoplasmic loop domain that protrudes far into the cytoplasm. Additionally, in the dark inactive state, the proximal region of the C-terminus (prior to residue D396, Figure 4A) takes on a conformation where, after the predicted 8^th^ helix, it is adjacent to the cytoplasmic extension of transmembrane helix 6, and close enough to form contacts. These contacts could potentially be ionic in nature, and our data support the importance of C-terminal phosphorylation sites, and Y382—which may also be phosphorylated—in phototransduction activation. Thus, C-terminal phosphorylation could provide the negative charge that is needed for the C-terminus to ionically interact with the arginine and lysine residue-rich 3^rd^ cytoplasmic loop, in the inactive state, prior to light-induced activation. Based on our homology modeling, we propose that upon photon-mediated conformational change to active-state melanopsin, the 5^th^ and 6^th^ transmembrane helices undergo an outward swing, moving the cytoplasmic ends of these helices—and the 3^rd^ cytoplasmic loop—away from the C-terminus. We also propose that the C-terminus, together with the 3^rd^ cytoplasmic loop, form a critical part of the G-protein binding pocket and may play a role in G-protein specificity or stabilization of the melanopsin—G-protein complex. Previous studies have also generated homology models of melanopsin utilizing squid rhodopsin (59, 60) or bovine rhodopsin (61), but these studies focused primarily on making computational predictions of melanopsin’s chromophore chemistry, and not focused on structural determinants of melanopsin—G-protein binding. However, these previous models also generated a predicted structure containing the features we highlight here (i.e. extended 5^th^ and 6^th^ transmembrane helices/3^rd^ intracellular loop and C-terminal conformation).

Our EPR spectroscopy measurements on melanopsin support the cytoplasmic extension of the 3^rd^ cytoplasmic loop because the spectral lineshape that is indicative of high spin-label rotational mobility at that region. Interestingly, while a clear difference was observed in full length and truncated melanopsin C268 EPR spectra, we did not observe dramatic differences in the spectral lineshape after light exposure in both samples. This result was unexpected, as we did predict to see light-induced conformational change reflected in the EPR spectra. Given the steric freedom at C268 and in the 3^rd^ intracellular loop region, it is possible that this region is unaffected by helical movement or conformational change due to its predicted distance from the membrane and from other structures. However, our model and data—including our EPR data of full length and C-terminal truncations—support the idea that the proximal C-terminus is adjacent to the 3^rd^ intracellular loop. Our model also predicts that upon light exposure, if melanopsin adopts a similar active conformation as the β_2_ adrenergic receptor, another family A GPCR, then we would expect these regions to separate and form the G-protein binding pocket. This would lead to the prediction that the EPR spectrum of the light-exposed full length melanopsin C268 sample should differ from the dark spectrum. It is possible that this movement happens as predicted by our modeling but does not produce a significant difference in the EPR spectral lineshape. Alternatively, it is also possible that the predicted interactions between these two regions, the C-terminus and the 3^rd^ cytoplasmic loop, remain intact even after light-induced activation. We observed no differences in the dark and light EPR spectra of the Δ365 truncated melanopsin C268 because this mutant lacks this C-terminal region. We therefore propose that our model of melanopsin activation is robust, but identification of structural and conformational changes of melanopsin after light exposure remains an important question for further study.

Precise structural and molecular determinants of GPCR selectivity to its cognate G-protein(s) are difficult to isolate, due the promiscuity of GPCRs for multiple G-proteins. While it would be erroneous to suggest that, like melanopsin, all Gαq-binding GPCRs require similar structural features (i.e. extended 5^th^ and 6^th^ transmembrane helices/3^rd^ cytoplasmic loop, positively charged cytoplasmic loops, and C-terminus contacting G-protein) for their selectivity, we propose that similar features are shared amongst rhabdomeric-type visual pigments. Of the available resolved protein structures of GPCRs, two are of rhabdomeric visual pigments: two squid rhodopsin structures and jumping spider rhodopsin (47, 48, 62). Similar structural features are observed in melanopsin’s predicted structure and the structures of the resolved rhabdomeric-type visual pigments. Specifically, the extended 5^th^ and 6^th^ transmembrane helices are a common feature amongst melanopsin and the invertebrate visual pigments, and like squid rhodopsin, melanopsin also has an extended C-terminus, much longer than vertebrate, ciliary-type image-forming visual pigments like rhodopsin. Melanopsin’s intracellular loops are predicted to possess a more uniform positive charge compared to the surface charges observed on the loops of squid and jumping spider rhodopsin, which are also positively-charged, but do possess regions of negative charge, particularly on the 5^th^ transmembrane portion of the 3^rd^ cytoplasmic loop. It is evident that this positive charge hold significance in melanopsin, but it remains unclear as to the extent of this feature’s significance for Gαq interaction. Given the enrichment of potential phosphorylation sites on melanopsin’s C-terminus and intracellular loops, this provides an interesting mechanism to modify the electrostatic potential of these regions, and thereby provides an interesting way to alter G-protein binding, and potentially specificity. Thus, with these findings, we identified critical regions that regulate melanopsin’s phototransduction activation to shed light onto ipRGC function and provide targets for protein engineering for use as an optogenetic tool or further study of its phototransduction.

## ACKNOWLEDGEMENTS

We thank and acknowledge NIH Training Grant T32 GM066706 awarded to J.C.V.L., generously made possible by Prof. Katherine Seley-Radtke and NIH R01 EY027202-01A1 awarded to P.R.R. We also thank Prof. Daniel D. Oprian for generously providing 1D4-antibody. M.P.D acknowledges support under the Cooperative Research Agreement between the University of Maryland and the National Institute of Standards and Technology Center for Nanoscale Science and Technology Award 70NANB10H193, through the University of Maryland.

NIST notes that certain commercial equipment, instruments, and materials are identified in this paper to specify an experimental procedure as completely as possible. In no case does the identification of particular equipment or materials imply a recommendation or endorsement by NIST, nor does it imply that the materials, instruments, or equipment are necessarily the best available for the purpose. The opinions expressed in this article are the authors’ own and do not necessarily represent the views of NIST.

## CONFLICT OF INTEREST

The authors declare that they have no conflicts of interest with the contents of this article.

## AUTHOR CONTRIBUTIONS

JCVL, STP, MPD, MG, and RJB conducted experiments. JCVL, EGC, JBW, VAS, and PRR intellectually contributed and conceived experiments. JCVL wrote the manuscript. PRR edited the manuscript.

---

**Supplemental Figure 1:** *Template sequence coverage of mouse melanopsin model*. (A & B) Mouse melanopsin homology model depicting amino acids on melanopsin that were modeled using amino acids on the *T. pacificus* template. Blue residues denote residues covered by template, pink residues are non-covered residues. (C) Template sequence coverage plotted on melanopsin’s amino acid sequence.

**Supplemental Figure 2:** *MATLAB/EasySpin simulation of experimental EPR data*. (**A**) Simulation spectrum fitting the full-length melanopsin C268 EPR spectrum or (**B**) melanopsin C268 Δ265, C-terminal truncated mutant. Rotational correlation times (τ_corr_) calculated from the simulation, depicted below the graph. Two components calculated for each construct, the 1^st^ and slower components represents a more immobile component, derived from spin-label attached to melanopsin, while the 2^nd^ and faster component represents a faster component, likely from excess, free spin-label in solution.

**Supplemental Figure 3:** Calcium imaging of double and triple melanopsin C-terminus proline point mutants. (**A**) Calcium imaging of HEK293 cells expressing melanopsin C-terminal mutants, synthesized with combinations of point mutations at residues H377, L380, and Y382 to proline residues. (**B**) Calculated activation rates of melanopsin constructs depicted in (A). All error bars represent S.E.M. of three independent transfections. Statistical significance tested by using Students t-test, *, **, ***, **** represent P-values <0.05, 0.01, 0.001, and 0.0001, respectively. All constructs compared to wild-type melanopsin’s rate, and statistical significance is indicated over individual bars.

**Supplemental Figure 4:** *Calcium imaging of melanopsin palmitoylation mutant*. Representative calcium imaging of melanopsin C364S, with mutation of melanopsin’s predicted C-terminus palmitoylation site, C364 to a serine residue. Error bars depict S.D.

**Supplemental Figure 5:** *2-dimensional schematics comparing mouse and armadillo melanopsin C-termini*.

